# Large errors increase the generalization of locomotor adaptation depending on the error direction

**DOI:** 10.1101/2022.06.01.494323

**Authors:** Digna de Kam, Wouter Staring, Dulce M. Mariscal, Gelsy Torres-Oviedo

**Author notes:** Corresponding author: Gelsy Torres-Oviedo, PhD, University of Pittsburgh, Department of Bioengineering, 4420 Bayard St, Suite 110, Pittsburgh, PA 15213, United States. Funding sources: This research was funded by an NSF CAREER grant to Gelsy Torres-Oviedo (1847891), Dulce Mariscal was funded by the NSF graduate fellowship program (NSF-GRFP 1747452).

## Abstract

Generalization in motor adaptation involves the transfer of movements beyond the adaptation context. We investigated the effect of size (large vs. small) and direction (unidirectional vs. bidirectional) of performance errors during adaptation on the generalization of walking patterns from a split-belt treadmill (training context) to overground (testing context). We hypothesized that unusual errors (i.e., large unidirectional or bidirectional errors) would serve as contextual cues limiting generalization. The size of unidirectional errors was modulated either implicitly (i.e., gradual vs. semi-abrupt split-belt perturbations) or explicitly (i.e., through instructed visual feedback). Bidirectional errors were induced by a sudden removal of the split-perturbation after a long adaptation period, resulting in errors in the opposite direction to those at the start of the adaptation period. Our findings did not support our hypothesis. We found that bidirectional, but not large, performance errors limited generalization across contexts, which could be mediated by two distinct mechanisms. On the one hand, bidirectional errors upon removal of the split-perturbation are also experienced when transitioning to overground walking. Thus, bidirectional errors may facilitate switching between distinct walking patterns, thereby limiting generalization. On the other hand, large unidirectional errors induce more motor adaptation, which might lead to more generalization.

## Introduction

Movement patterns that are adapted through interactions with the world can be generalized from trained to untrained conditions. For instance, arm movements that are modified when reaching in one direction influence reaches when aiming at other directions^1–3^. Similarly, walking patterns adapted on a treadmill partially carry over to walking overground^4,5^. Generalization of movements from trained to untrained contexts is limited because of contextual cues such as sensory information^6^ or body state^7^ linking motor memories to the context in which they were learned. It is important to understand contextual cues that regulate the generalization of movements from trained to untrained contexts because the effectiveness of rehabilitation interventions relies on the transfer of movements learned in the therapy setting to daily life situations.

Performance errors in the training environment are a contextual cue that regulates generalization of adapted movements. This might be because large performance errors laying outside subjects’ ordinary range are attributed to the training context, rather than subjects’ faulty movements^4^. Thus, reducing the size of performance errors during adaptation may be beneficial for generalization of movements across contexts. Performance errors can be reduced using different strategies. For instance, they can be reduced through implicit processes in which subjects adapt their motor commands without being aware of it^1,8^. Alternatively, they can be reduced through explicit strategies based on cognitive constructs of the task^9–11^. It has been shown that cognitive processes can influence the generalization of locomotor movements across contexts^5,12^, raising the question of whether reducing errors through implicit or explicit strategies would have a differential effect on generalization.

In addition to error size, the direction of performance errors may also regulate the generalization of adapted motor patterns. Specifically, performance errors during a locomotor adaptation task on a split-belt treadmill can be unidirectional (e.g., negative step length asymmetry) or bidirectional (e.g., negative and positive step length asymmetries). Unidirectional performance errors are observed upon experiencing the unusual split-belt environment, in which the legs move at different speeds^13^. Bidirectional performance errors occur when, in addition to the initial errors, subjects experience errors in the opposite direction upon removal of the split-environment, which has been adopted as the “new normal”^14^. Thus, removing the split-environment is equivalent to experiencing a disturbance in the opposite direction. Bidirectional errors are inherently present in generalization studies in which the split environment is removed multiple times to assess and compare adaptation effects (i.e., aftereffects) across different contexts (e.g. aftereffects on treadmill vs. aftereffects overground). However, little is known about the impact of bidirectional errors on generalization. We speculate that the unusual nature of bidirectional errors could be used as a contextual cue^6,15^, and thereby, reduce the generalization of movements.

We investigated the effect of error direction (i.e., unidirectional vs. bidirectional step length asymmetry) and error size (i.e., small vs. large step length asymmetry magnitude) on the generalization of locomotor adaptation from the treadmill (training context) to overground (testing context). We hypothesized that unusual errors (i.e., large or bidirectional errors) would serve as contextual cues limiting generalization. To test this, we assessed the impact of reducing unidirectional errors, either implicitly or explicitly, on the generalization of movements (Experiment 1). Moreover, we investigated the effect of error direction on generalization and the extent to which this effect is modulated by error size. (Experiment 2).

## Results

### Reducing performance errors through explicit or implicit strategies limits the generalization of locomotor patterns across contexts

In ***Experiment 1***, we assessed the effect of performance error size on the generalization of locomotor patterns across contexts. We used step length asymmetry as our metric of performance errors because it is a conventional measure that robustly characterizes the adaptation of gait on split-belt walking^16^. We were particularly interested if performance errors (i.e., step length asymmetry) reduced through implicit vs explicit strategies yielded comparable effects on generalization. To this end, we compared a group that was designed to exhibit large performance errors during adaptation during split-belt walking (Abrupt^UNI^) to a group that reduced performance errors using explicit (Feedback^UNI^) or implicit (Gradual^UNI^) strategies (Figure 1). The large error group experienced large performance errors due to a semi-abrupt introduction of the split-perturbation (40 stride ramp). A similar perturbation was used for a Feedback^UNI^ group that was instructed to maintain symmetric step length using visual feedback. Performance errors were reduced using an implicit strategy in the Gradual^UNI^ group, which was exposed to a gradual split-perturbation (600 stride ramp). Figure 2A shows the performance errors during split-belt walking for each of the three groups, which significantly differed in the maximal performance error shortly after encountering the full split-perturbation (Oneway ANOVA MaxError: F(2,28)=9.3, p=0.001,η^2^=0.41). Post-hoc tests revealed that the MaxError was significantly larger in the Abrupt^UNI^ compared to the Feedback^UNI^ (p=0.001) and Gradual^UNI^ (p=0.023) groups. Moreover, MaxErrors were comparable between the Feedback^UNI^ and Gradual^UNI^ groups (p=0.36). Performance errors in the steady state of split-belt walking also differed across groups (Oneway ANOVA lateError: F(2,28)=5.1, p=0.013, η^2^=0.27). Specifically, lateErrors were larger in the Gradual^UNI^ compared to the Feedback^UNI^ groups (p=0.011). No significant differences in lateError between the Abrupt^UNI^ vs Feedback^UNI^ (p=0.55) and between the Abrupt^UNI^ vs the Gradual^UNI^ groups (p=0.11).

**Figure 1:**
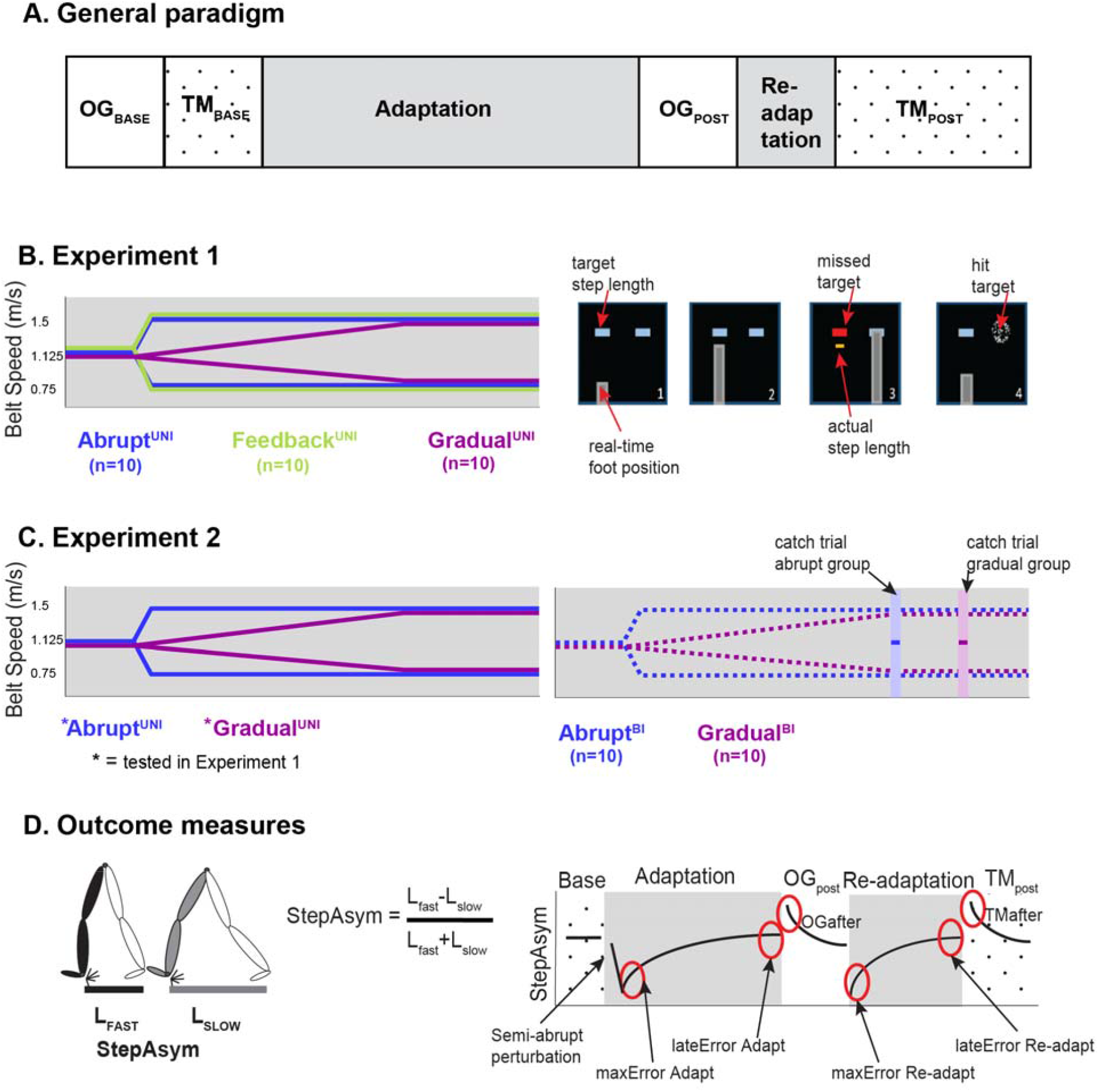
Methods. **A: Experimental conditions.** Subjects first performed baseline walking overground (OG_base_) and on the treadmill (TM_base_) before they were subjected to the split-environment (Adaptation). Subsequently, they walked overground (OG_post_) to assess generalization of movements across contexts. Participants in the Abrupt and Feedback groups were re-adapted on the treadmill, such that aftereffects in the trained context could be assessed (TM_post_). The adaptation trial was varied across the groups to determine the effect of error size and direction on generalization. **B: Experiment 1**. Left panel shows the belt speed profiles for subjects in the Abrupt^UNI^ (large error), Feedback^UNI^ (small error explicit correction), and Gradual^UNI^ groups (small error implicit correction). Participants in the Feedback^UNI^ group were presented with symmetrical step length targets (right panels) as well as real time foot position. They were also informed on their actual step length relative to the target. **C: Experiment 2**. Belt speed profiles for the unidirectional (left panel) and bidirectional error groups (right panel). Note that the unidirectional error groups were the same subjects tested under experiment 1. **D: Outcome measures**. Step length asymmetry (left panel) was used to quantify performance errors and aftereffects. Performance errors were quantified after subjects reached the full split condition (maxError) and at the end of (re-)adaptation (lateError). Aftereffects were quantified in the untrained context (OG_post_) for all groups and also in the trained context (TM_post_) in groups with an abrupt perturbation.

**Figure 2:**
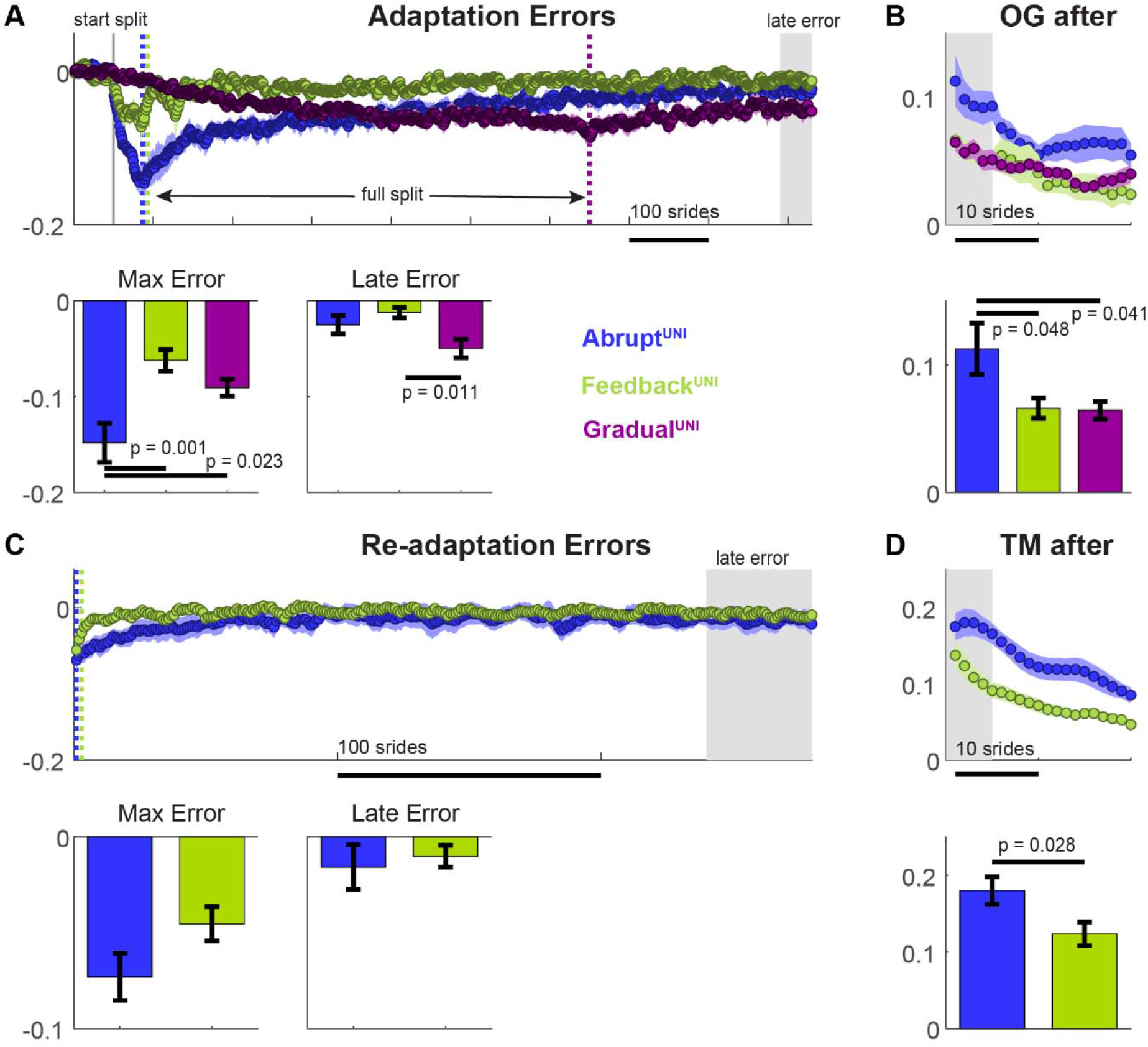
Experiment 1. **A: Adaptation errors.** Time courses of errors during the adaptation period (top panel) and performance error outcomes (bottom panels). The vertical grey line indicates the start of the split perturbation. Colored dotted lines indicate the instant were the full split condition was reached for each group. **B: Overground aftereffects**. Time courses (upper panel) and averages (bottom panel) of step length asymmetry aftereffects during overground walking following the Adaptation period. **C: Re-adaptation errors**. Time courses of errors during the re-adaptation period (top panel) and performance error outcomes (bottom panels). C**: Treadmill aftereffects**. Time courses (upper panel) and averages (bottom panel) of step length asymmetry aftereffects during treadmill walking following the re-adaptation period. * Group means and standard error of the means are presented in all panels. The grey box indicates the time window used to caclulate the late error (Adaptation and Re-adaptation) and the aftereffects (OG after and TM after). All time were smoothed for visual purposes using a moving average window of 5 strides.

We evaluated the effect of manipulating the error size implicitly or explicitly on the generalization of walking patterns by comparing across groups (Abrupt^UNI^, Feedback^UNI^, Gradual^UNI^) the aftereffects of step length asymmetry when walking overground. We found that generalization was limited in the small compared to the large performance error groups. (Figure 2B). Specifically, a oneway ANOVA yielded a significant effect of group for overground aftereffects (F(2,28)=4.3, p=0.024, η^2^=0.24). These aftereffects were significantly larger in the Abrupt^UNI^ group compared to the Feedback^UNI^ (p=0.048) and Gradual^UNI^ groups (p=0.041). The Feedback^UNI^ and Gradual^UNI^ groups exhibited comparable overground aftereffects (p=0.997). In sum, reducing performance errors during split-belt adaptation implicitly or explicitly limits the generalization of motor patterns to the overground context.

### Reduced performance errors during split-belt walking result in smaller aftereffects in the training context

Participants in the Abrupt^UNI^ and Feedback^UNI^ group were re-adapted to the split-condition to assess the effect of error size during split-belt walking on aftereffects in the training (i.e. treadmill) context (Figure 2C). During the readaptation period, these groups did not significantly differ in their performance errors (maxError: t(18)=1.83, p=0.09; lateError: t(18)=0.44, p=0.67). Despite this similarity, the subsequent aftereffects on the treadmill were smaller in the Feedback^UNI^ compared to the Abrupt^UNI^ group (t(18)=2.38, p=0.028, d=1.07; Figure 2D). Thus, our results show that smaller performance errors upon the first exposure to split-belt walking resulted in reduced aftereffects in both the training (treadmill) and the testing context (overground).

### Bidirectional performance errors reduce the generalization of locomotor patterns only when the initial performance errors are large

In ***Experiment 2***, we assessed the effect of error direction (bidirectional vs unidirectional performance errors) during split-belt walking on the generalization of locomotor patterns across contexts. To this end we compared participants with unidirectional performance errors (Experiment 1) to participants that experienced additional performance errors in the direction opposite to the initial ones. Opposite errors were elicited by means of a catch trial, involving 10 strides of tied-belt walking, after a long period (i.e., at least 150 strides) of split-belt walking at the full speed difference (Figure 1). We further argued that bidirectional errors could affect generalization differently in the case of large (Abrupt) vs small (Gradual) initial performance errors (i.e. interaction between error size and direction). Thus, Experiment 2 was designed as a 2 by 2 experiment with error size and error direction as independent variables of interest, resulting in 4 experimental groups (Abrupt^UNI^, Gradual^UNI^, Abrupt^BI^ and Gradual^BI^).

Figure 3A illustrates performance errors of each of the abrupt and gradual groups during adaptation. Consistent with Experiment 1, we found that performance errors after introduction of the initial split-perturbation were larger in the Abrupt than Gradual groups (MaxError: F_ErrorSize_(1,36)=17.6, p=0.0002; 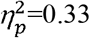). On the other hand, the MaxError was not different between groups with unidirectional vs bidirectional errors (F_ErrorDirection_(1,36)=0.71, p=0.41; F_ErrorSize*ErrorDirection_(1,36)=0.30, p=0.59). Moreover, performance errors at the end of the split-perturbation (lateError) were not different across groups (Error Size: F_ErrorSize_(1,36)=2.6, p=0.12; Error Direction F_ErrorDirection_(1,36)=0.78, p=0.38; F_ErrorSize*ErrorDirection_(1,36)= 0.69,p=0.41). Participants in the bidirectional groups also exhibited performance errors in the direction opposite to the initial perturbation during the catch trial. These errors were not different between the Abrupt^BI^ and Gradual^BI^ groups (t(18)=0.30, p=0.77).

**Figure 3:**
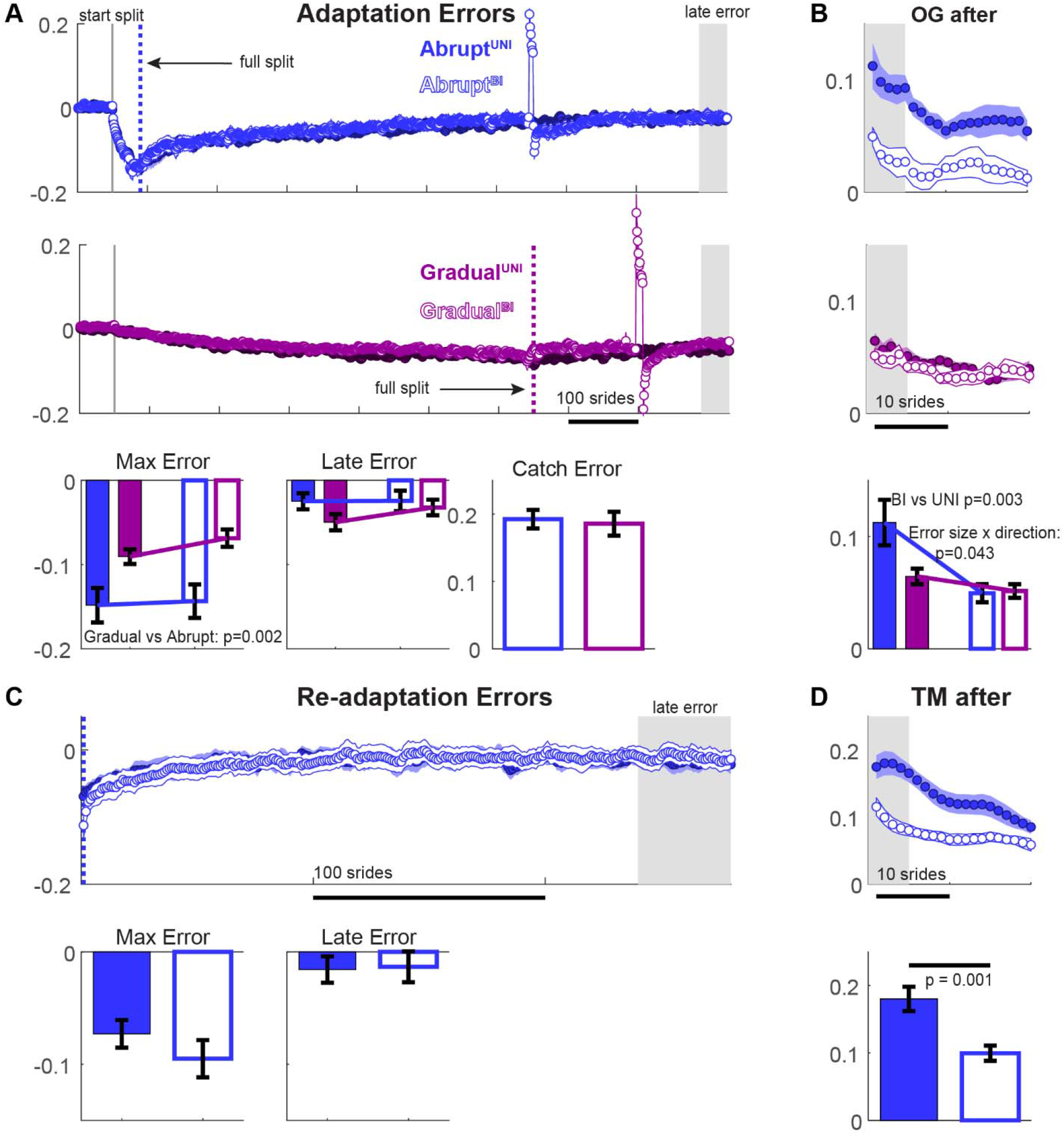
Experiment 2. **A: Adaptation errors.** Time courses of errors during the adaptation period (top panel) and performance error outcomes (bottom panels). The vertical grey line indicates the start of the split perturbation. Colored dotted lines indicate the instant were the full split condition was reached for each group. **B: Overground aftereffects**. Time courses (upper panel) and averages (bottom panel) of step length asymmetry aftereffects during overground walking following the Adaptation period. **C: Re-adaptation errors**. Time courses of errors during the re-adaptation period (top panel) and performance error outcomes (bottom panels). C**: Treadmill aftereffects**. Time courses (upper panel) and averages (bottom panel) of step length asymmetry aftereffects during treadmill walking following the re-adaptation period. * Group means and standard error of the means are presented in all panels. The grey box indicates the time window used to calculate the late error (Adaptation and Re-adaptation) and the aftereffects (OG after and TM after). All time courses (except for the catch trial) were smoothed for visual purposes using a moving average window of 5 strides.

We found that overground aftereffects were regulated by both error direction and size (Figure 3B). Specifically, the ANOVA yielded a significant main effect of error direction (F_ErrorDirection_(1,36)=10,3, p=0.003, 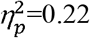), due to smaller aftereffects in subjects who had been subjected to bidirectional compared to unidirectional performance errors. While no significant main effect of Error Size was observed (F_ErrorSize_(1,36)=3.8, p=0.06), a significant interaction was present (F_ErrorSize*ErrorDirection_(1,36)=4.4, p=0.043, 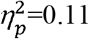). The interaction effect resulted from a large reduction of overground aftereffects following adaptation with bidirectional vs unidirectional errors in the abrupt groups, but not in the gradual ones. Thus, our results show that bidirectional errors limit the generalization of locomotor patterns across contexts, but this effect is mediated by the size of the errors upon introduction of the split-perturbation.

### Bidirectional performance errors during split-belt walking result in smaller aftereffects in the training context

We investigated whether the observed reduction in aftereffects upon bidirectional performance errors was unique to the overground context. To this end, aftereffects on the treadmill were compared between the Abrupt^UNI^ and Abrupt^BI^ groups after they were first re-adapted to the split-condition. We found that readaptation errors were comparable between the groups (MaxError: t(18)=1.07, p=0.30; lateError: t(18)=-0.13, p=0.90; Figure 3C). On the other hand, subsequent aftereffects on the treadmill were significantly smaller in the Abrupt^BI^ vs Abrupt^UNI^ group (t(18)=3.79, p=0.001 d=1.69; Figure 3D). Thus, our results show that bidirectional performance errors during adaptation reduced aftereffects in both the training and testing context.

## Discussion

Our study aimed at investigating the effect of performance error size (large vs. small) and direction (unidirectional vs. bidirectional) on the generalization of locomotor adaptation from the treadmill (training context) to overground (testing context). The size of unidirectional performance errors was modulated either implicitly (i.e., gradual split-belt perturbation) or explicitly (i.e., through instructed visual feedback). We found that reducing performance errors, either implicitly or explicitly, led to small aftereffects on the treadmill (training context) and overground (testing context). Thus, small error sizes limit the generalization of adapted motor patterns likely because they induce less adaptation. Moreover, bidirectional errors also led to small aftereffects in both contexts (i.e., treadmill and overground), likely because bidirectional errors were generated by suddenly removing the split-perturbation, which is conventionally used for assessing aftereffects on the treadmill or overground. Thus, the repeated exposure to removing the split-perturbation might facilitate switching between distinct walking patterns within the training environment or across distinct walking environments.

The generalization of movements across contexts is limited by specific cues that may promote the linking of motor patterns to the context in which they were adapted^6,15,17^. For example, visual information unique to the training context promotes context-specificity of motor patterns in reaching tasks^6,15^. In locomotor adaptation on a split-belt treadmill, removing unique visual information from both the training (treadmill) and testing context (overground) facilitates the generalization of locomotor patterns^17^. Similar to visual cues, it was proposed that performance errors during adaptation may serve as a contextual cue that impacts the generalization of adapted motor patterns^4^. Specifically, it was suggested that subjects attribute errors outside their ordinary range to the training context (treadmill), rather than to their own performance, thereby limiting the generalization of adapted motor patterns beyond the training context^4^. Consistently, we hypothesized that large errors, as well as bidirectional errors, would enhance the context-specificity of adapted motor patterns, thereby limiting generalization across contexts (i.e. from treadmill to overground). However, our findings do not support this hypothesis.

Specifically, in Experiment 1 we found that small, rather than large (out of the ordinary), performance errors diminished generalization. Reducing initial performance errors using implicit or explicit corrections resulted in smaller aftereffects overground (testing context), which contrasts with prior findings^4,18^. It must be noted that in these previous studies participants experienced bidirectional errors, whereas we only induced large unidirectional errors, possibly explaining the discrepancy in our results. We also found that aftereffects on the treadmill (training context) were smaller if subjects used an explicit strategy (visual feedback) to reduce their errors, which is at odds with previous findings^9,19^. The discrepancy between our results and previous findings may have resulted from the difference in the initial large performance errors across our studies. Specifically, we eliminated the large initial performance errors that are typically observed during split-belt paradigms by using a semi-abrupt perturbation which enabled participants to fully counteract the split-belt perturbation through explicit corrections. Conversely, participants in the study by Roemmich et al., 2016^9^ exhibited large initial performance errors because they experienced an abrupt split-belt perturbation that could not be fully corrected with an explicit strategy. We suggest that these large initial performance errors may serve as teaching signals that drive locomotor adaptation. Consequently, reducing the size of initial performance errors may allow for less adaptation of motor patterns, thereby also limiting their generalization across contexts.

Moreover, the results from Experiment 2 do not support the hypothesis that the unusual nature of bidirectional errors would serve as a contextual cue linking the adapted motor patterns to the treadmill context. According to this hypothesis, aftereffects would be small in the testing context (overground) but not in the training context (treadmill). In contrast, we found that bidirectional errors reduced aftereffects in both contexts. We believe that bidirectional errors led to small aftereffects because a similar environmental transition (i.e., removal of the split-perturbation) is used to originate bidirectional errors during the catch trial and to measure aftereffects during post-adaptation. Such repeated exposure to a specific environmental transition (i.e. perturbation introduction or removal) has shown to result in more efficient switching between motor patterns^20,21^. Thus, we suggest that generalization of movements across contexts was reduced by bidirectional errors because they facilitate switching between motor patterns upon removal of the split-perturbation. Interestingly, the effect of bidirectional errors on generalization appeared to be modulated by the size of the initial errors. Specifically, bidirectional errors only reduced generalization in the case of large initial errors (Abrupt groups). We believe that this is due to a floor effect in the gradual groups, who exhibited already little generalization following unidirectional errors. In sum, bidirectional errors limit the generalization of motor patterns across context and this effect is mediated by the size of initial performance errors. This finding is important to take into account in the interpretation and design of generalization studies that inherently involve bidirectional errors to assess aftereffects in the trained context^4,18^.

In sum, our findings suggest that performance errors out of subjects’ ordinary range do not promote linking of the motor pattern to the context in which they were adapted. Instead, errors larger than usual enhance generalization because they allow for more adaptation. Moreover, bidirectional errors facilitate switching between motor patterns upon environmental transitions, thereby limiting generalization. We found that reducing the magnitude of performance errors using either implicit or explicit strategies limited the generalization of adapted motor patterns across contexts to the same extent. Nevertheless, it is plausible that (partially) distinct mechanisms underlie these reductions in generalization due to the different neural circuitry involved in implicit and explicit motor adaptation. Specifically, implicit motor adaptation is a cerebellar-dependent process^22–27^ and involves recalibration of forward internal models of the environment based on sensory prediction errors^28,29^. These errors were smaller in our gradual paradigm, likely resulting in less recalibration and thus smaller aftereffects overground. In contrast to the cerebellar-driven implicit adaptation, explicit error corrections involve a more conscious strategy hinting at a stronger involvement of cerebral structures in motor adaptation^30,31^, possibly through connections with the cerebellum^32,33^. Overground aftereffects in the visual feedback group, which reduced errors through explicit corrections, were likely smaller because participants no longer used the cognitive strategy learned on the treadmill during overground walking. Specifically, the use of explicit corrections depends on the presence of visual feedback^9,34^, which was not provided during overground walking. Moreover, the transition from treadmill to overground walking may serve as a contextual cue^6,15^ that makes participants aware of the perturbation removal. Hence, they may stop using the explicit correction strategy developed on the treadmill. In sum, explicit error corrections limit the carryover of movements from the treadmill to overground walking, resulting in smaller overground aftereffects.

While implicit and explicit error correction strategies may impact generalization through different neural mechanisms, it must be noted that implicit and explicit adaptation are not completely independent. Specifically, implicit motor adaptation still occurs if subjects use an explicit strategy to correct their performance errors^8,9,19,35^. Nevertheless, our visual feedback most likely increased the relative contribution of the explicit component to counteract the perturbations as compared to the gradual group whose corrections are mostly implicit^36^. Taken together, reducing performance errors using implicit and explicit correction strategies limit generalization to the same extent, but the underlying mechanisms are partially distinct.

Our study has a few limitations that need to be considered. First, in order to accurately measure our primary outcome for generalization (overground aftereffects) in Experiment 1, we tested aftereffects on the treadmill (training context) after subjects had been readapted to the split-environment. This may have had an impact on the aftereffects in the training context, which can be considered a study limitation. Specifically, repeated exposure to environmental transitions allows subjects to switch motor patterns more quickly^20,21^, and thus, treadmill aftereffects in our study were likely underestimated due to prior exposure to perturbation removal. A second limitation is that our experimental design did not allow for a reliable comparison of treadmill aftereffects in the Gradual vs Feedback groups. Hence, future studies are needed to provide more insight in potential differences in context-specificity of adapted motor patterns after implicit vs explicit error corrections.

Our findings have important clinical implications. First, in rehabilitation programs, the generalization of trained motor patterns to daily life situations is essential. For example, training on a split-belt treadmill can be used to reduce gait asymmetry during overground walking in individuals with a stroke^37–39^. However, some stroke survivors show a limited generalization of trained motor patterns from the treadmill to overground walking^39^, perhaps because their motor system is less sensitive to smaller step length asymmetry errors^40^. Therefore, it is possible that generalization of training effects in clinical populations can be enhanced by exposure to larger performance errors during training. Second, our findings suggest that repeated exposure to environmental transitions results in more effective switching between motor patterns (i.e. smaller aftereffects). The ability to flexibly switch between motor patterns is key in daily life situations in which we are often exposed to sudden perturbations. We suggest that training protocols aimed at improving motor flexibility will likely benefit from including sudden environmental changes.

In conclusion, large initial performance errors upon an exposure to an environmental transition facilitate generalization of adapted motor patterns across contexts, likely because they induce more adaptation. On the other hand, bidirectional errors during a motor adaptation task limit generalization, because it promotes flexible switching between motor patterns upon sudden environmental transitions. This has important implications for the design of motor adaptation experiments in which bidirectional errors are commonly used.

## Methods

### Subjects

A group of neurologically intact young individuals was included in this study (n=60, 24 males, age=25.6±4.7). To be eligible for participation, subjects had to: 1) have no prior experience to split-belt walking; 2) have no orthopedic conditions interfering with the assessment 3) have no neurological conditions; 4) have no contra-indications for performing moderate intensity exercises; 5) use no medication that could affect cognitive or motor functions. All participants provided written informed consent prior to study participation. The study was approved by the University of Pittsburgh Institutional Review Board and was conducted in accordance with the Declaration of Helsinki.

### Locomotor paradigm

#### General paradigm

The experiment involved participants walking both overground and on a treadmill. The treadmill was a split-belt treadmill with two separate belts that were driven by independently controlled motors. A thin plastic divider was placed in the middle of the treadmill (between the legs) to ensure that each foot was on its separate belt. For safety requirements participants wore a harness and they were able to hold on to a side rail if necessary. The harness did not support body weight during walking. The subjects were informed of the starting and stopping of the treadmill. The treadmill was stopped between each of the testing trials for a brief period. Subjects were instructed to look straight ahead while walking. The general protocol common to experiments 1 and 2 is shown in Fig. 1A. Specifically, the protocol consisted of: 1) six minutes of over ground walking on a 10 meter walkway to capture subject’s normal overground walking pattern; 2) baseline walking on the treadmill at different speeds (slow: 100 strides at 0.75 m/s; fast: 100 strides at 1.5 m/s; medium: 450 strides at 1.125 m/s) to familiarize subjects with the treadmill and capture their normal treadmill walking pattern; 5) adaptation to the split-environment (900 strides evolving towards a 2:1 belt-speed ratio, 0.75 and 1.5 m/s); 6) six minutes of overground walking to capture aftereffects in the untrained context (i.e. generalization); 7) a second adaptation trial (300 strides) to bring subjects back to their adapted state and 8) a post-adaptation period on the treadmill (600 strides) to capture subjects aftereffects in the trained context. The belt speed profiles for the first adaptation period were systematically varied between the groups to determine the effects of error size and direction on the generalization of motor patterns from the treadmill to overground walking. Lastly, subjects were instructed to stand still between the end of the adaptation and beginning of post-adaptation trials because stepping in place or moving their feet could potentially reduce their aftereffects.

#### Experiment 1

Our study was designed to determine the effect of error size and error direction on generalization of walking patterns across contexts. The focus of *Experiment 1* was on error size, thereby, error direction was kept constant across groups (i.e. unidirectional errors only). We specifically investigated whether reducing the size of performance errors would have a distinct effect on generalization if these errors were corrected by implicit vs. explicit strategies. To this end, we contrasted a group that was designed to exhibit large performance errors to two groups that reduced their performance errors using either implicit or explicit strategies. The large error group (Abrupt^UNI^, Fig 1B, blue trace) experienced large performance errors due to a semi-abrupt introduction of the split-perturbation. A similar perturbation was used for a group that used an explicit strategy to reduce the error size using visual feedback (Feedback^UNI^, Fig. 1B, green trace). Specifically, the split-belt condition (i.e., speed difference between the feet) was introduced over the course of 40 strides. After the 40 strides ramp period subjects walked in the split-environment (i.e. 2:1 belt-speed ratio) for 860 strides. The ramp period was chosen such that it would allow subjects in the Feedback^UNI^ group to correct their errors as the speed difference was introduced, which is not possible with a full abrupt split-perturbation^9,19,35^. While this semi-abrupt perturbation leads to smaller initial errors than a full abrupt perturbation (i.e., 2:1 ration introduced abruptly), which is conventionally used in large error groups ^4^, this distinction does not have an impact on generalization. Namely, we verified that the semi-abrupt group and a full abrupt group (n=10) had similar generalization of motor patterns from the treadmill to overground walking (0.10 ± 0.05 vs 0.11 ± 0.06; p=0.74). Thus, the Abrupt^UNI^ group characterizes well the generalization of a group with large errors.

The Feedback^UNI^ group was instructed to maintain symmetric step lengths using visual feedback (Fig. 1B, right panels). Step length targets for both legs represented each individual’s average step length of both legs obtained during baseline walking at 1.125 m/s. Each time the participant initiated a step, a gray bar representing the inter-ankle distance was displayed on the screen until the instant of heel strike. Participants were instructed to hit the targets with the bar by regulating their step length (i.e., inter-ankle distance at heel strike). The step length targets had a tolerance of ±5 centimeters such that deviations of < 5 cm of the intended step lengths were allowed. Targets “exploded” if subjects reached them successfully, whereas they turned red upon failure to reach them. In addition, we displayed a yellow horizontal bar, in failed steps, to provide feedback for the missing distance between the participant’s step length and the target step length. Subjects in the Feedback^UNI^ group underwent a brief training trial of 150 strides to familiarize with the feedback.

Lastly, the belt speeds’ profile during the adaptation period for the Gradual^UNI^ group was designed to reduce performance errors in an implicit manner^4^. This was done by gradually changing the belt speeds from tied to split over the course of 600 strides, followed by a hold period of 300 strides at the 2:1 belt-speed ratio. We chose this long ramp period because it reduces subjects’ awareness of the stride by stride change in belt speeds (Mariscal et al. 2020). Note that the second adaptation trial, consisting of a full abrupt split-perturbation induces large performance errors, particularly in subjects that are naive to an abrupt perturbation (Gradual group). Given that those large errors may confound the subsequent treadmill aftereffects, the treadmill post-adaptation trial was not considered subjects in the gradual group.

#### Experiment 2

In *Experiment 2*, we examined the effect of error direction (i.e., unidirectional vs. bidirectional errors) on the generalization of movements across contexts. Moreover, we investigated if the effect of error direction on generalization would be modulated by error size (i.e., small vs. large errors). We speculated that bidirectional errors would limit generalization, particularly in combination with large errors, because this would yield more errors outside subject’s ordinary range. Thus, we conducted a 2 by 2 design in which both error size and error direction were systematically varied across groups (Abrupt^UNI^, Abrupt^BI^, Gradual^UNI^, Gradual^BI^; see Fig 1C. for adaptation protocols). In the unidirectional error groups, subjects experienced errors in only one direction during the adaptation period in the training context. In the bidirectional errors groups, subjects experienced errors in one direction when the split-perturbation was introduced and in the opposite direction, compared to those initially experienced, when the split-perturbation was briefly removed during the adaptation period ^4^. To this end, the treadmill was stopped during the adaptation period and subjects performed a catch trial that involved 10 strides of tied-belt walking at 1.125 m/s; after which the treadmill was stopped again and resumed split-belt walking at a 2:1 belt speed ratio until the end of the adaptation period. Error size was varied across groups by using a semi-abrupt (large errors) vs. a gradual introduction of the split-perturbation (small errors). Note that the adaptation protocols for the Abrupt^UNI^ and Gradual^UNI^ groups were similar in *Experiment 1*. Therefore, data of subjects in those groups were used for both *Experiment 1 and 2*.

For the bidirectional error groups, perturbation removal during adaptation was conducted in subject’s adapted state, because only in that case the split-perturbation is experienced as the “new normal”, such that the removal of it elicits large errors in the opposite direction^4,14^. Thus, in the Abrupt^BI^ group, the split-perturbation was removed after 600 strides of adaptation (i.e. 560 strides of exposure to the full-split condition), whereas in the Gradual^BI^ the split-pert was removed after 750 strides (i.e. 150 strides of exposure to the full-split condition).

### Data collection and processing

Data were collected using a 14-camera VICON motion analysis system. Reflective markers were placed bilaterally on the following landmarks. The foot (metatarsal 1 and calcaneus), the ankle (lateral malleolus), the knee (lateral femoral epicondyle), the hip (greater trochanter), and the pelvis (anterior and posterior iliac spine). Kinematic data were collected at 100 Hz. During treadmill walking, ground reaction forces were recorded from each leg separately with a frequency of 2000 Hz. For treadmill walking, we used ground reaction force data to detect gait events. Specifically, we determined the instants at which the feet landed (i.e., heel-strike: Fz>10N) or were lifted from the ground (i.e. toe-off: Fz<10N). For overground walking, we used kinematic data for event detection similar to prior studies^4,5^.

### Data analysis

#### Kinematic parameters

Step length asymmetry (StepAsym) was used to quantify the size of performance errors as well as aftereffects^4^ (Figure 1D). Specifically, we defined StepAsym as the difference between consecutive steps of the legs in terms of step length, where step length was defined as the distance between the leading and the trailing limb ankles at heel strike. In our definition, StepAsym is positive when the step length of the fast leg (i.e. dominant) is larger than the one of the slow leg (non-dominant). StepAsym was expressed in units of distance and normalized to the total stride length in order to account for differences in step sizes across subjects^5^ (Equation 1).

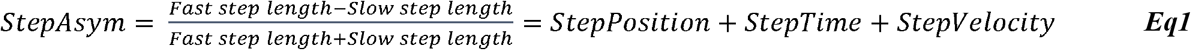

#### Outcome measures

We characterized subjects’ errors (stepAsym) during the first and second adaptation trials in terms of 1) maximum error size, 2) late error size. Specifically, the maximum error size (maxError) was quantified as the maxim error (i.e. most negative stepAsym) during the first 15 strides after subjects experienced the full split-perturbation (i.e. at the end of the ramp period). This maximum was determined using a running average of 5 strides. We computed errors in the steady state of adaptation (lateError) by averaging stepAsym over the last 40 strides of the adaptation trials. For groups with bidirectional errors we also computed errors during the catch trial (average of the first 5 strides).

We quantified after-effects in the untrained (overground) and trained context (treadmill) to respectively assess generalization and adaptation of motor patterns. To this end, we computed the average values of stepAsym, over the first 5 strides of the post-adaptation trials (OG-post and TM-post). Kinematic outcome measures were corrected for baseline biases by subtracting the median values of subjects’ baseline trial at medium speed (i.e. last 40 strides). This baseline correction was done for overground and treadmill trials separately. Lastly, we discarded the very first 1 and the last 5 strides of treadmill trials to avoid any effects of starting and stopping of the treadmill.

#### Statistical analysis

For experiment 1, outcome measures for the first adaptation and overground post-adaptation trials were compared between the Abrupt^UNI^, Feedback^UNI^ and Gradual^UNI^ groups using a oneway ANOVA. In the case of a significant main effect of group, we performed Tukey corrected post-hoc tests for group comparisons. Outcome measures for the second adaptation and treadmill post-adaptation trials were considered for the Abrupt^UNI^ and Feedback^UNI^ groups only. For these outcome measures, differences between the groups were assessed using an independent samples t-test.

For experiment 2, we assessed the effect of error direction and error size on overground aftereffects. To this end, we performed a 2-way ANOVA with error direction (UNI and BI) and error size (Abrupt and Gradual) as independent factors. We also included an interaction term (Error Direction * Error Size) to assess if the effect of error direction was modulated by error size. Moreover, we assessed the effect of error direction on aftereffects in the trained (treadmill context) for the Abrupt groups only using an independent samples T-test.

We reported P-values, F-values, and t-values for all group analyses. In the case of a significant group effect, effect sizes were reported as well (η^2^ for one-way ANOVAs, 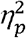 for two-way ANOVAs and Cohen’s d for independent samples t-tests). An alpha-level of 0.05 was used for all statistical tests.

## Author contributions

Experiment design: GTO, DK, WS, DM. Data collection: WS, DK, DM. Data analysis: WS, DK, DM. Writing manuscript: DK, GTO

## Data availability statement

Data files are available from the authors upon request.

